# Intrinsically disordered N-terminal and structured DNA-binding domains jointly regulate progesterone receptor transcriptional condensates

**DOI:** 10.64898/2026.04.21.719910

**Authors:** Louna Raluy, Dominik Saul, Catalina Romero-Aristizabal, Borja Requena, Lara I. de Llobet Cucalon, Robyn Laura Kosinsky, Nadine Youssef, Carla Garcia-Cabau, Maciej Lewenstein, Xavier Salvatella, Gorka Muñoz-Gil, Maria F. Garcia-Parajo, Juan A. Torreno-Pina, Priyanka Sharma

**Affiliations:** Institut de Pharmacologie et de Biologie Structurale, IPBS, Université de Toulouse, CNRS, UPS, Toulouse, France; Division of Endocrinology, Mayo Clinic, Rochester, MN 55905, USA; Robert and Arlene Kogod Center on Aging, Mayo Clinic, Rochester, MN 55905, USA; Robert Bosch Center for Tumor Diseases, Stuttgart, Germany; Centre de Regulació Genomica (CRG), The Barcelona Institute of Science and Technology (BIST), Dr. Aiguader 88, Barcelona, Spain; ICFO-Institut de Ciències Fotòniques, The Barcelona Institute for Science and Technology (BIST), 08860 Barcelona, Spain; Institute for Research in Biomedicine (IRB Barcelona), The Barcelona Institute of Science and Technology, Barcelona, Spain; ICREA (Institució Catalana de Recerca i Estudis Avançats), Pg. Lluís Companys 23, 08010 Barcelona, Spain; University of Innsbruck, Department for Theoretical Physics, Technikerstr. 21a, A-6020 Innsbruck, Austria; Present address: Department of Macromolecular Structure, Centro Nacional de Biotecnologia-CSIC, C/ Darwin, 3, 28049 Cantoblanco Madrid, Spain

## Abstract

Transcriptional condensates (TCs) are dynamic, membrane-less assemblies of proteins and nucleic acids that form at specific genomic loci to control gene expression. The primary drivers of TC formation and function are long stretches of polypeptide chains that lack well-defined tertiary structure, known as the intrinsically disordered regions (IDRs) of Transcription Factors (TFs). However, the role of well-structured domains such as the DNA-binding domain (DBD) in condensate formation and downstream regulation remains poorly understood. Here, we investigated whether the DBD of the Progesterone Receptor (PR), a model TF regulated by hormone binding, contributes to the formation of PR TCs. By combining Single Particle Tracking (SPT) with deep-learning-based single-trajectory analysis, we demonstrated that both the IDR and the DBD are required for the formation of PR condensates. Furthermore, transcriptomic analysis showed that mutations of either of the two domains result in the expression of a distinct set of genes, with functional assays corroborating these findings and linking domain-specific transcription to differential effects on cell proliferation and migration. Together, our results show that domains beyond IDRs also play a role in TC formation and function and uncover domain-specific regulation of transcriptional and oncogenic programs.

## Introduction

Cells exploit compartmentalization as a key regulatory mechanism to control non-equilibrium biochemical processes. While classic examples involve lipid-membrane-bound organelles, recent studies have proposed biomolecular condensation as a general principle of membrane-less compartmentalization (1). The resulting assemblies, known as biomolecular condensates, can transiently and specifically recruit biomolecules, enabling the spatial and temporal organization of biochemical reactions, signal transduction, and transcriptional regulation in the cell. Importantly, this dynamic organization allows cells to respond rapidly to environmental or signaling cues that regulate the stability of condensates. The aberrant biomolecular condensation of specific biomolecules due to mutations, mis-splicing, or overexpression has also been associated with different diseases and, in cancer, with the dysregulation of gene expression, the disruption of cellular homeostasis, and oncogenesis (2–5). Understanding how aberrant condensation plays a role in cancer can therefore highlight new opportunities for targeted therapeutic interventions.

Transcription factors (TFs) are nuclear proteins that regulate the initiation and maintenance of gene expression. Over the past decade, it has become clear that TF activity can arise through their capacity to assemble into high-order species, perhaps by undergoing a phase transition, leading to the formation of clusters or transcriptional condensates (TCs). Such TCs can then facilitate the recruitment of components of the transcriptional machinery, including RNA Polymerase II and the mediator complex (6–12). In addition to this important activity, TCs can also locally reorganize chromatin enhancer hubs, further enabling a precise control of transcription plasticity in cancer (13–19). From a molecular and a mechanistic perspective, it is well-established that the intrinsically disordered regions (IDR) of TFs play a central role in their ability to (I) cluster or condense and (II) recruit interacting molecules (12, 20–24). In contrast, the role of the structured DNA-binding domain (DBD), despite noteworthy *in vitro* work using ensemble measurements, has been less explored (25).

Nuclear receptors (NR) constitute a family of TFs that exclusively trigger transcription only upon exposure to specific steroid hormones (26–28). This tunability makes them ideal candidates to investigate how TCs form and contribute to transcriptional activity in a time-resolved fashion. Indeed, we and others have recently shown that the progesterone receptor (PR), the estrogen receptor (ER), the glucocorticoid receptor (GR), and the androgen receptor (AR) are all able to form condensates only upon binding to their relevant hormone (8, 10, 23, 29–33). While some of these studies have focused on the intrinsically disordered N-terminal domain (NTD) of NRs, which plays a key role in TC formation and transactivation, little is known about the role of the structured DBD in controlling the formation and maintenance of TCs. Most importantly, whether and how the IDR and the DBD synergistically control the properties of TCs and the transcriptional program of NRs remains unknown.

Here, we addressed these unresolved questions by using PR as a model system and by applying Single Particle Tracking (SPT), a powerful single-molecule-based method that enables the study of individual TFs and their interactions with biologically plausible components such as chromatin in a minimally invasive manner in living cells (22, 31, 33, 34). In addition, several SPT-associated data analysis methods have been proposed to infer different subpopulations based on the lateral diffusion of TFs. One prominent example is the “Spot-On” analysis, which is based on the fitting of lateral displacements using a kinetic mixture model and allows distinguishing between subpopulations of molecules (e.g., “bound” vs “free”) (35). Recently, machine learning (ML) has also extended the possibilities of classifying diffusive behaviors of TFs using short trajectories. Indeed, using ML, we previously identified two different diffusing subpopulations of PR and, furthermore showed that the low mobility population corresponded to PR molecules diffusing within a condensate (29).

By relying on distinct PR mutants combined with SPT and different SPT analyses, including machine learning, we here unveiled the role of the structured DBD along with the intrinsically disordered NTD in the formation and regulation of TCs. Bulk transcriptomics and functional experiments further revealed that mutations in these two domains result in the expression of strikingly distinct sets of genes, both linked to oncogenic pathways. Taken together, our results provide additional evidence that domains beyond the IDRs contribute to biological condensate formation and function.

## Results

### Both the NTD and the DBD control the slowly diffusing population of PR

To examine how the NTD and the DBD control the spatiotemporal behavior of PR in living cells at the single molecule level, we generated two PR mutants in living MCF7 breast cancer cells: (I) one lacking the complete intrinsically disordered N-terminal domain of the PR (“ΔNTD”) and (II) another with point mutations in the DNA binding region of PR that obliterate its affinity for DNA (for simplicity, we will refer to this mutant as “ΔDBD”) (**Fig. 1A**). To avoid any side interactions between the mutants and the untagged endogenously expressed PR, we used a Knockdown and Rescue (pKAR) system where the endogenous receptor is knocked down using RNA interference and instead expresses a rescue RNAi-resistant PR (**Supplementary Fig. 1A-B**) (36).

**Figure 1:**
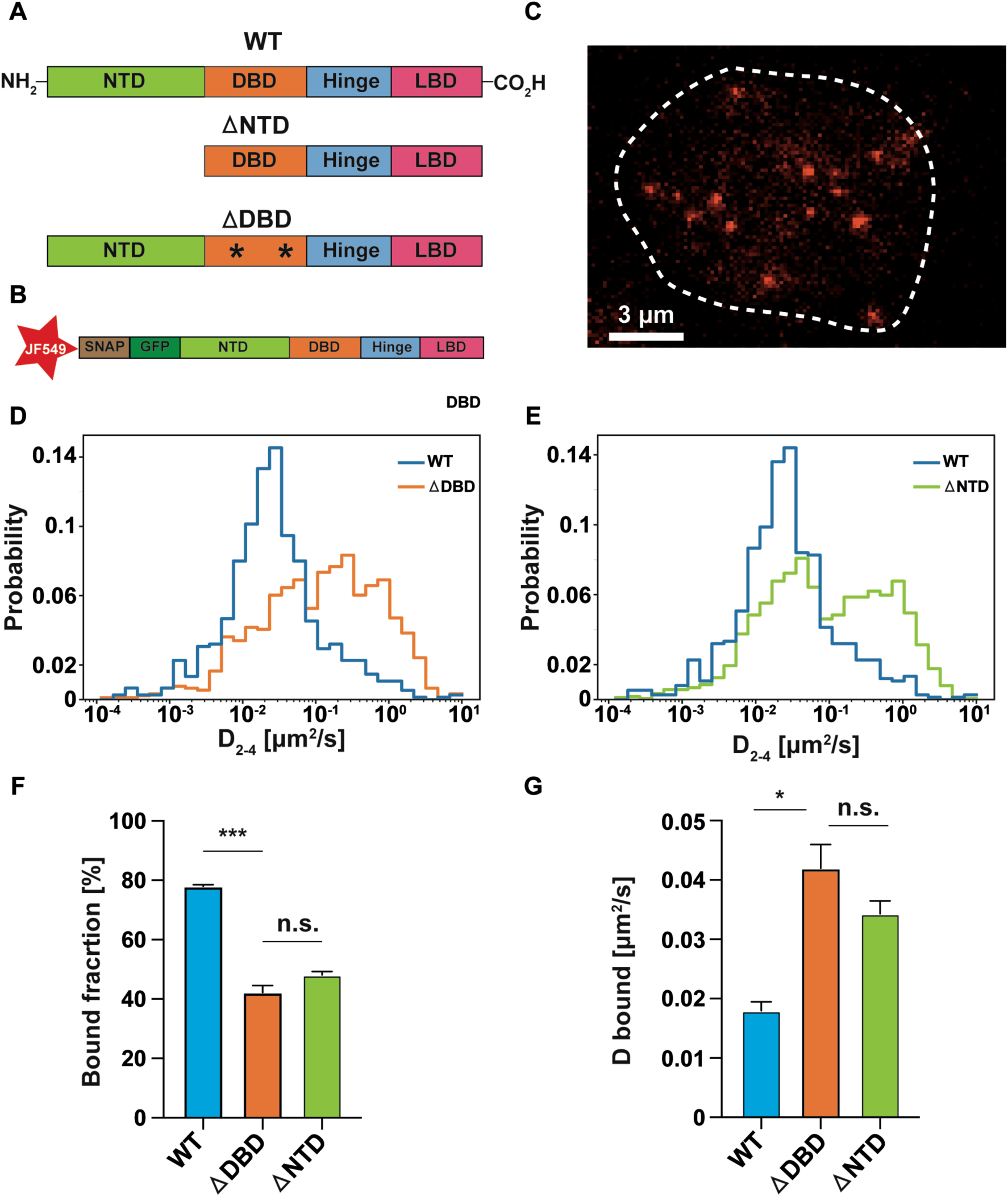
The lateral diffusion of WT and mutants (βDBD or βNTD) in the nucleus of living breast cancer cells. **(A)** Schematic representation of the different constructs (WT, βDBD, and βNTD) used throughout the study. (**B**) Schematic representation of the construct used for SPT in the nucleus. A SNAP-tag was introduced in the N-terminal domain of the GFP-PR. (**C**) Representative frame of the SPT video. Individual PR molecules labelled with Janelia Fluor 549 are visualized using red color coding in the nucleus (white outline) of MCF7 cells. (**D-E**) Normalized histograms of the instantaneous diffusion coefficients (D_2-4_) for hormone-exposed WT, βDBD, and βNTD extracted from the MSD curves for each single SPT trajectory. (**F-G**) Bound fraction and diffusion coefficients from the three constructs extracted from the Spot-On analysis. The error bars correspond to the standard error of the mean. Results of a one-way analysis of variance (ANOVA) test are shown as: n.s. for not significant, * for p-value< 0.05, and *** for p-value<0.001. SPT data is extracted from at least 2 independent experiments, 8 cells, and 750 trajectories per condition.

To enable SPT, we designed PR constructs in which we introduced a SNAP tag at the N-terminus of WT PR, βNTD, or βDBD (**Fig. 1A-B**) (29, 37). We then performed SPT using single-molecule sensitive highly inclined thin illumination (HILO) of SNAP-Janelia Fluor 549 tagged PR molecules in the cell nucleus (**Fig. 1C**). SPT videos were acquired after incubating the cells (WT, βDBD, or βNTD) with the PR ligand, a synthetic progesterone derivative R5020, at 10nM for 60 min. These experimental conditions were used since, in our previous report, they were shown to induce PR condensate formation (29). In order to visualize fast dynamic events such as DNA-binding, we recorded the SPT videos with a framerate of 15 ms. After identifying the centroid position of each single PR molecule with a localization precision of ∼25 nm, we reconnected each position as a function of time to obtain individual trajectories that report on the diffusion of PR and its mutants.

We first analyzed the SPT trajectories by extracting the instantaneous diffusion coefficient (D_2-4_) by linearly fitting the 2^nd^-4^th^ points of the mean-square displacement (MSD) curve for each trajectory and generated distributions of all the D_2-4_ values from hundreds of trajectories from multiple cells (**Fig. 1D-E**, **Supplementary Fig. 1C**). Our results showed an increase of D_2-4_ values for both mutants compared to WT, with median values of D_2-4_ = 0.024 μm^2^/s (WT), 0.132 μm^2^/s (ΔDBD), and 0.078 μm^2^/s (βNTD). Moreover, compared to the WT histogram, which largely shows a single distribution centered around low D values, both mutants exhibit binomial distributions, suggesting the existence of different mobility sub-populations, in particular, the emergence of an increased diffusing population. Thus, these results reveal that compared to WT, both the βDBD and βNTD mutants exhibited increased diffusion, indicating that both the DBD and the NTD influence the low mobility of PR molecules.

To extract quantitative information on these apparently different subpopulations in the lateral diffusion of PR and the mutants, we applied the “Spot-On” analysis. In our case, the data was best fitted to a two-population model, which we further considered as “free” and “bound” sub-populations (35). Remarkably, the bound fraction extracted for WT is 77.9±0.6%, which is significantly reduced to 42.3±2.3% and 48.1±1.1 % for βDBD and βNTD, respectively (**Fig. 1F**, **Supplementary Fig. 1D**). Moreover, the reduced bound sub-population of the PR mutants (βDBD or βNTD) showed an increased mobility compared to WT, as indicated by the diffusion coefficients of the bound fraction: 0.018 ± 0.001 μm^2^/s, 0.042±0.004 μm^2^/s, and 0.034 ± 0.002 μm^2^/s for WT, βDBD, and βNTD, respectively (**Fig. 1G**). Altogether, D_2-4_ and Spot-On analysis indicate that both the DBD and the NTD control the low-mobility population and the bound fraction of PR.

### Deep learning single-trajectory analysis identifies the NTD and DBD-dependent PR diffusion within condensates

The relatively short duration of our SPT trajectories represents a challenge in characterizing the dynamics of the PR. To gain further resolution in the available data, we use STEP, a deep-learning-based method to predict the anomalous diffusion exponent and diffusion coefficient for every frame of an input trajectory and extract segments of constant diffusive properties (38) (**Supplementary Fig. 2**). We used the value of anomalous exponent *α* (see Materials and Methods for details) as a measure of diffusion behavior, with *α=*1 corresponding to Brownian diffusion and *α <* 1 to anomalous diffusion (*i.e.* subdiffusion). We generated 2D histograms of the anomalous exponents and diffusion coefficients for each of the detected segments of WT and the two mutants (βDBD and βNTD). In the case of the WT, the 2D histogram shows one main cluster centered around D values of 0.1 μm^2^/s and exhibiting *α* values below 0.5 (**Fig. 2A**), which we previously identified as a diffusion within a PR condensate (29). In contrast, the two PR-mutants show a shift toward higher D and *α* values, concomitant with an overall reduction in the fraction of the low-mobility component. To further uncover these changes in the distributions, we computed the difference between WT and βDBD /βNTD probability distribution functions across the plane (*α*, D), showing a remarkable shift in the distributions toward higher values in both variables (**Fig. 2B**).

**Figure 2:**
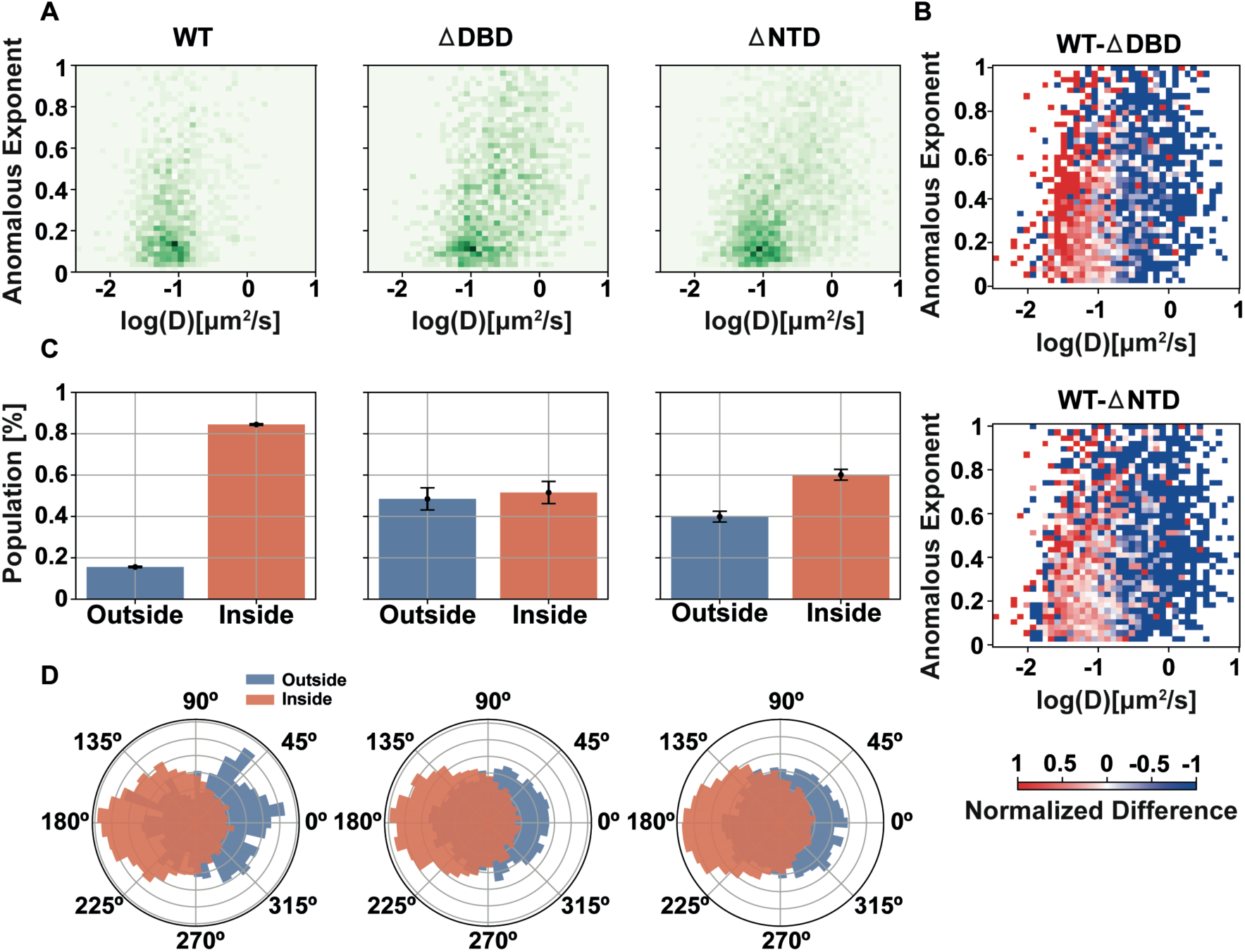
Deep learning analysis (STEP) reveals two sub-populations within TCs. (**A**) Histograms of the predicted diffusion coefficient and anomalous exponent for every segment of WT, βDBD, and βNTD found by means of STEP and a kernel changepoint detection algorithm. (**B**) Difference between the parameter distributions of the different constructs, calculated from the normalized joint histograms as shown in A. (**C**) Percentage of segments per population as extracted from a k-means clustering algorithm. (**D**) Angle distribution between consecutive segments for each sub-population and construct. The data used for STEP analysis has been extracted from at least 2 independent experiments, 8 cells, and 903 trajectories per condition.

Next, we clustered the segments of each distribution by means of the k-clustering algorithm and obtained two different clusters that we associate with diffusion inside and outside of condensates (**Supplementary Fig. 3**) (29). By computing the percentage of segments in each population, we obtained that WT contains 84.5±0.3% and 15.5±0.3% of diffusion inside and outside of condensates, respectively (**Fig. 2C**). Importantly, ΔDBD (51.5±5.3%) and ΔNTD (60.1±2.6%) show a remarkable reduction in the population that diffuse inside condensates, indicating that both protein domains contribute to the formation of PR condensates.

Finally, we computed the angles between successive steps of each classified segment of the two populations and the three constructs. We plotted the distribution of the angles for each population, showing that while the population of diffusion outside of condensates has an isotropic angle distribution, the population of diffusion within condensates displays an expected increase at 180^0^ (**Fig. 2D**) (29). Taken together, STEP identifies a (I) population of PR molecules diffusing within condensates and (II) that this population is strongly dependent on both the DBD and the NTD.

### NTD and DBD distinctly regulate the transcription programs

Our results so far indicate that the NTD and the DBD contribute similarly to the capacity of the PR to form condensates, but do not provide information on how they influence the transcriptional activity of the receptor. To this end, we performed mRNA-sequencing in cells expressing WT and the two PR mutants (βDBD or βNTD) under progestin exposure. A principal component analysis (PCA) of mRNA-sequencing data showed a distinct clustering of the results obtained for WT, βDBD, and βNTD before and after R5020 induction (**Supplementary Fig. 4**). Upon progestin treatment, we identified 2,293 differentially expressed genes (DEGs) in cells expressing WT, 2,418 DEGs in cells expressing βNTD, and only 1,165 DEGs in ΔDBD cells, reflecting a markedly attenuated response to progestin. In total, this analysis identified 3,291 DEGs, including 773 DEGs (23.5%) that were found in all three cases (WT, βDBD and βNTD) (**Fig. 3A**). In addition, WT and ΔNTD shared additional 839 DEGs, bringing the total overlap between the two to 1,612 (52%) (**Fig. 3A**). Similar comparison between WT and βDBD cells yielded 77 additional DEGs for a total overlap of 850 (32.6%), and 123 DEGs when comparing βNTD and βDBD cells, for a total of 896 DEGs (33.3%) (**Fig. 3A**). Furthermore, 192 genes (16.5% of the total 1,165 DEGs) were uniquely regulated in ΔDBD cells, while a larger set of 683 genes (28.2% of 2,418 DEGs) was uniquely regulated in the ΔNTD cells, suggesting a more prominent role for the NTD in fine-tuning transcriptional outputs. Moreover, 604 genes (26.3% of 2,293 DEGs) of unique genes were expressed in WT cells (**Fig. 3A**). Overall, these results indicate that inactivating DNA binding results in more profound changes in the transcriptional response to progestin and suggest that βNTD and βDBD may affect a distinct set of cellular processes.

**Figure 3:**
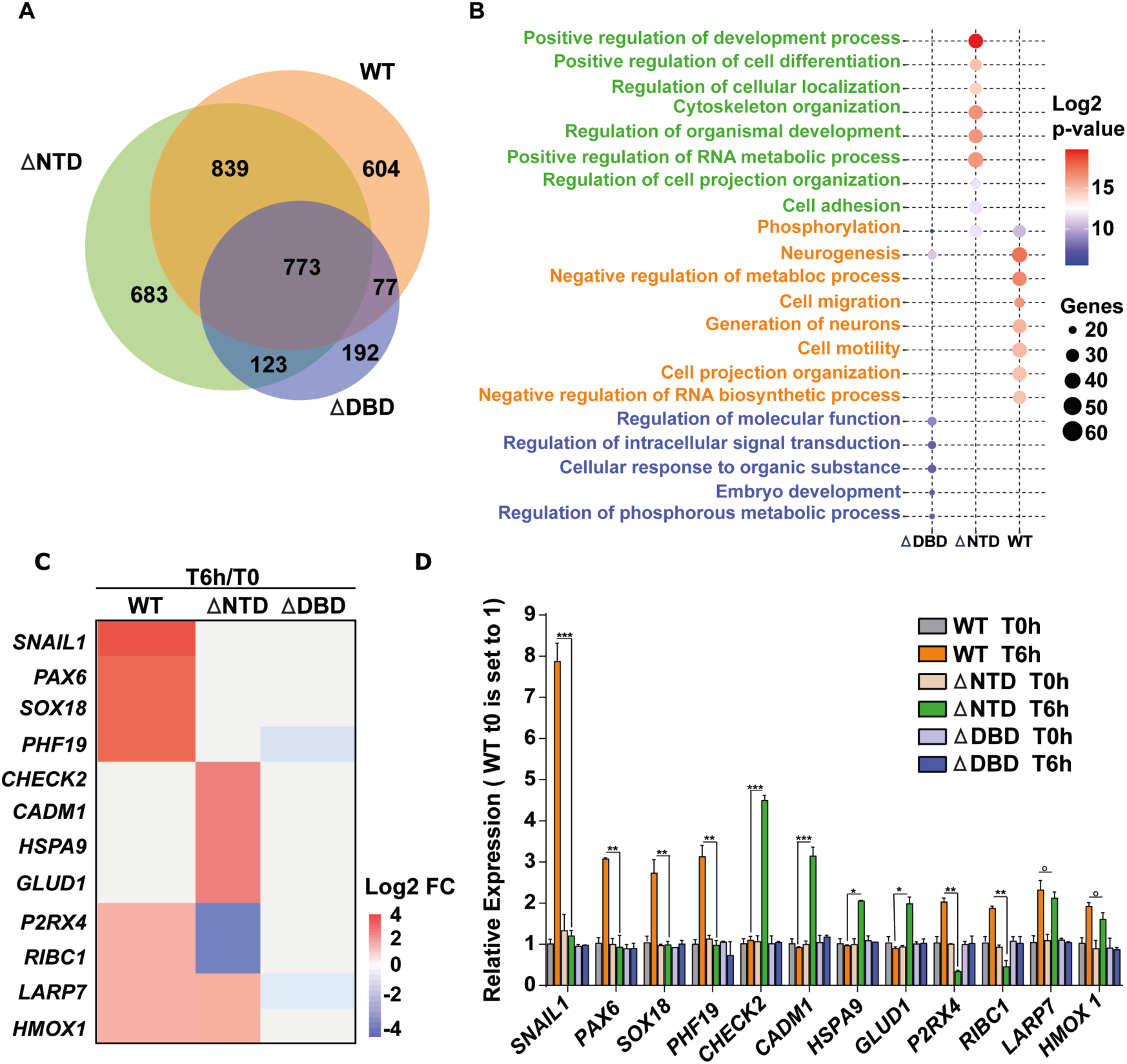
Progestin-induced transcriptional profiling reveals distinct regulatory gene programs for individual domains of the PR. (**A**). Venn diagram representing the overlap of differentially expressed genes (Fold change > 1.5 or <1/1.5 and p value < 0.05) modulated by R5020 induction (10nM for 6 hours) in MCF7 cells specifically expressing the WT and mutants (βDBD and βNTD). (**B**). Representative plots of key gene sets for R5020-induced differentially expressed genes in WT, βDBD, and βNTD expressing MCF7 cells. Log2 p-values are indicated in color and size, showing the number of enriched genes. (**C**). Heatmap representing the log2 fold change in the expression of the representative gene set of “progestin-responsive genes” from the RNA-sequencing analysis performed in WT, βNTD, or βDBD MCF7 cells. (**D**) Quantitative RT-PCR validation in MCF7 cells expressing WT or mutants in the absence or presence of R5020. Changes in mRNA levels were normalized to GAPDH mRNA. Data represented as WT, as set to 1, mean ± SEM of four biological experiments.

To get a more granular view of processes differentially affected by the two mutants, we performed Gene Ontology analysis (**Fig. 3B**). The genes uniquely affected in ΔNTD cells were significantly enriched in developmental as well as fundamental cellular processes, such as cell adhesion, cellular localization, and RNA metabolic pathways, which are frequently deregulated during oncogenesis. This suggests that the NTD is critical for the optimal expression of PR-dependent genes that help maintain proper cellular programs. In comparison, genes uniquely regulated in ΔDBD cells were enriched for molecular functions related to signal transduction and metabolic processes. Meanwhile, genes uniquely regulated in WT cells were enriched for cellular processes, such as cell migration, cell motility, cell organization, and negative regulation of metabolic processes and pathways, which were largely absent from both mutants. These findings suggest that mutations in either of the domains (NTD and DBD) regulate a distinct set of cellular processes via controlling the expression levels of unique sets of genes. Moreover, disruption of either the NTD or the DBD compromises PR’s ability to drive expression of genes associated with oncogenic cell migration in MCF7, a breast cancer cell line, highlighting that both domains play a role in PR-mediated tumorigenic programs.

To further validate the transcriptional differences in cells expressing WT and the two mutants, and examine their potential participation in tumorigenic programs, we selected a subset of representative genes that are both differentially expressed (**Fig. 3C**) and associated with key tumor-related processes (39–50). This panel included genes involved in epithelial-mesenchymal transition (*SNAIL1*), developmental regulation (*PAX6*, *SOX18*), chromatin remodeling (*PHF19*), DNA damage response (*CHEK2*), cell adhesion (*CADM1*), stress response (*HSPA9*, *HMOX1*), cellular metabolism (*GLUD1*), purinergic signaling (*P2RX4*), and transcriptional regulation (*RIBC1*, *LARP7*) (39–50). We validated the differential gene expression using RT-qPCR (**Fig. 3D**). Remarkably, *SNAIL1*, *PAX6*, *SOX18*, and *PHF19* were uniquely up-regulated only in WT cells upon progestin-induction (**Fig. 3D**). In contrast, *CHECK2*, *CADM1*, *HSPA9*, and *GLUD1* were only uniquely up-regulated in βNTD cells, highlighting a distinct transcriptional response associated with deletion of the PR activation domain (**Fig. 3D**). Additionally, *P2RX4* and *RIBC1* were down-regulated in βNTD cells, while both genes were up-regulated in WT cells. Progestin-induced expression of *LARP7* and *HMOX1* remained unchanged between WT and ΔNTD cells. Importantly, none of these genes showed significant changes in expression in ΔDBD cells. Collectively, these observations highlight the potential of the NTD domain to fine-tune transcription regulation to maintain the condensate landscape for key oncogenic processes.

### The NTD and DBD domains are required for PR-driven oncogenic processes

Our DEGs, as well as SPT analysis, indicate that both the NTD and the DBD are required for the PR TCs formation and regulate distinct cellular processes. To functionally validate the results, we examined phenotypes of MCF7 breast cancer cells expressing WT, ΔNTD, or ΔDBD. Because many PR-regulated genes disrupted in the domain-depleted mutants (ΔNTD or ΔDBD) are linked to cell proliferation, we first assessed progestin-induced proliferation in MCF7 cells expressing WT, ΔNTD, or ΔDBD. Progestin induction significantly increased the proliferation in WT cells, whereas this effect was significantly reduced in both ΔNTD and ΔDBD cells (**Fig. 4A**). Consistent with the gene expression followed by the Gene Ontology enrichment analysis, we then evaluated progestin-stimulated cell migration using a transwell migration assay (48). MCF7 cells expressing ΔNTD or ΔDBD showed markedly impaired migration compared to WT (**Fig. 4B**, *left*; **Supplementary Fig. 5**) and by acetic acid–based extraction (**Fig. 4B**, *right*). Together, these results demonstrate that both the NTD and the DBD of PR are required for the establishment of PR-dependent transcriptional programs that promote oncogenic processes such as cell proliferation and migration.

**Figure 4:**
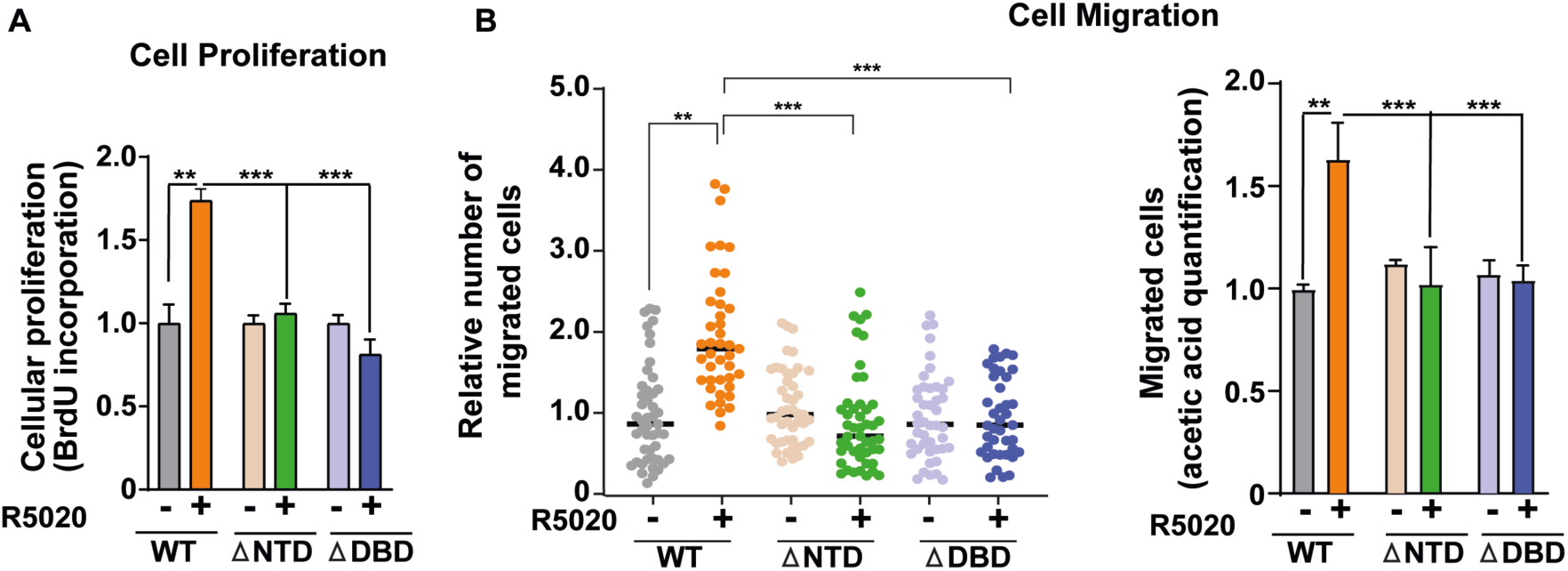
Both protein domains regulate oncogenic processes. (**A**) Cell proliferation of MCF7 cells expressing the WT or the two mutants before and after the induction of R5020. Data represent median ± SEM of at least eight biological experiments. (**B**). Cell migration of MCF7 cells expressing WT or mutants (βNTD or βDBD) before and after the induction of R5020. Data represent median ± SEM of at least three biological experiments. (*Left*) Dot plot showing the relative number of migrated cells in the indicated condition. (*Right*) Bar plot representing the acetic acid quantification of the migrated cells. *p value < 0.05; **p value < 0.01; ***p value < 0.01.

## Discussion

Here, we have presented a single-molecule biophysical study dissecting the contribution of distinct PR protein domains to transcriptional condensate formation in cancer cells. SPT combined with trajectory analysis revealed reduced low-mobility populations and a diminished bound fraction in both βNTD and βDBD mutants, as extracted from MSD and Spot-On analysis, respectively. STEP, a deep-learning SPT analysis pipeline, revealed a two-population-based behavior of PR in the nucleus. Based on the mobility and anomalous exponent clustering by STEP, we identified the anomalous diffusion and the low mobility population as the population belonging to the diffusion within condensates (29). Importantly, condensate formation was strongly dependent on both the NTD and DBD, indicating that these domains regulate condensate assembly and stability.

We also assessed the contribution of PR protein domains on global gene expression by performing bulk mRNA-sequencing. Remarkably, breast cancer cells expressing βNTD or βDBD-mutants displayed markedly distinct transcriptional profiles, with only a partial subset of genes shared between the two mutants, underscoring domain-specific regulation of transcription. Notably, the NTD mutant led to a significant effect on the transcription activity of the genes associated with oncogenic processes. In contrast, the DBD mutant resulted in the transcription alteration of the genes associated with metabolic pathways and cellular signaling functions. These transcriptional findings were validated by functional assays of cell migration and proliferation, confirming that disruption of either domain compromises PR-dependent cellular programs.

Based on our SPT and global gene transcription and functional assays, we propose a novel regulatory mechanism of TCs. It has been suggested that the concerted action between DBDs and IDRs might regulate gene transcription (51, 52). However, how this regulatory interplay might act in living cells remains to be further studied. In recent years, with the application of live cell imaging, the interaction between IDR domains, via possible biocondensation, and interacting partners is emerging as a complementary mechanism for TF selectivity. Indeed, using single-molecule imaging, a recent paper suggests that the interaction between scaffolded unstructured domains of TFs, rather than DNA binding, accounts for TF specificity (53). Importantly, in this study, they did not observe evidences for condensate formation.

In the case of PR, our previous study established that the PR forms condensates (29). Here, we compared Spot-On together with STEP and obtained similar populations of low and higher mobility in WT and in both mutants, suggesting that both types of analysis are retrieving the same biophysical mechanism, namely PR condensates. Importantly, when the NTD is depleted, there is still a 60.1% fraction of PR molecules diffusing within a condensate, suggesting that the structured DBD is a main regulator of condensate formation. However, the DNA binding specificity might be compromised by PR-dependent IDR-IDR interactions as suggested in the above-mentioned study (53). In the case of the DBD mutant, 51.5% of PR molecules still diffuse within a condensate, showcasing the canonical contribution of IDR in regulating condensate formation. In this case, and since there is no DNA-binding by PR, PR-dependent DEGs are uniquely mediated by PR-mediated IDR interactions with other TFs and related transcription machinery. Further studies would be required to address the (I) DNA-binding sites and (II) IDR-IDR interactions promoted by aberrant PR condensates. Overall, the differential gene expression profile and the SPT data point to an IDR and DNA-binding dependent mechanism in mediating functional PR transcriptional condensates in breast cancer.

Altogether and to the best of our knowledge, our work provides the first systematic and comprehensive analysis of the distinct roles of TF domains in regulating condensate formation and transcription program. While numerous studies have highlighted the contribution of TF IDRs to TCs, often using artificial gene arrays or focusing solely on IDR manipulation, we establish a tunable condensate model controlled by hormone stimulation in breast cancer cells. By examining both the DBD and the NTD of PR, we uncover a prominent role for the structured DBD in condensate formation and demonstrate how each domain shapes transcriptional output. Based on our observations, we propose that IDRs and structured DBDs jointly affect TCs formation and function, opening a new paradigm in the field of biocondensates.

## Materials and Methods

### Plasmids

The original plasmid expresses the PR isoform-B under a tetracycline-controllable promoter (TetOff system, Clontech) and a SNAP-tag at the N-terminal domain of the GFP (29). Taking this plasmid as the backbone, we introduced a shRNA-based mutation, thereby knocking down endogenous PR expression. We engineered the wild-type (WT) PR and mutants by either mutating the DBD-PR and deleting the NTD-PR (referred to as βDBD or βNTD in the manuscript).

### Cloning

The original plasmid (pGFP-PRB plasmid) expresses the PR isoform-B under a tetracycline-controllable promoter (TetOff system, Clontech) and a SNAP-tag at the N-terminal domain of the GFP (29). This plasmid was used to insert the shRNA sequence (GGGACAACATAATTATTTGTG) within the exon 3 of the PR-B coding gene, and was confirmed by sequencing. The inhibition activity of the inserted shRNAs sequence was confirmed by the reduced level of PR-B (**Supplementary Fig. 1A**). To generate the catalytically inactive DBD domain (ΔDBD) of the pGFP-PRB plasmid, the potential set of amino acids (G585 to E585, S586 to G586, C587 to A587, V589 to A589, A604 toT604, R606 toW606, R637 to A637, K638 to A638) were mutated (54–56). Whereas, to generate the ΔNTD, the gene fragment (Integrated DNA Technologies) was synthesized by digestion with Bsu36 (R0524S, NEB) and PstI-HF (R3140S, NEB) restriction enzymes, followed by purification and ligation using T4 DNA ligase (M0202S). The final plasmid was confirmed by Sanger sequencing. To generate the ΔNTD pGFP-PRB plasmid, the pGFP-PRB shRNA1 plasmid was cut with BsrGI (R3575S, NEB), Bsu36I (R0524S, NEB), and SrfI (R0629S, NEB) restriction enzymes. After gel purification (QIAquick Gel Extraction Kit), the 5827 and 1044bp fragments were preserved. The following linkers were cut with BsrGI (R3575S, NEB) and Bsu36I (R0524S, NEB) with prior annealing of the 2 oligos (Integrated DNA Technologies) with Buffer 4 from NEB at 95°C for 5 minutes:

> Linker: 5’-CGAGCTGTACAAGTCCGGACTCAGATCTCGAGCGTTGCCTGAGGCCGGAT-3’ 5’-ATCCGGCCTCAGGCAACGCTCGAGATCTGAGTCCGGACTTGTACAGCTCG-3’

Next, a gene fragment without the NTD domain (Integrated DNA Technologies) was digested with BglII (R0144S, NEB) and PstI-HF (R3140S, NEB), ligated with T4 DNA ligase (M0202S). The final plasmid was also confirmed by Sanger sequencing.

### Cell culture and electroporation

MCF7 Tet-off cells (Clontech, Cat No. 631154) were grown on Dulbecco Modified Eagle Medium (DMEM) high-glucose media supplemented with 10% Tet-free Fetal Bovine Serum, 2mM L-glutamine, 1 mM sodium pyruvate, and 100 U/ml penicillin-streptomycin. Cells were electroporated simultaneously with the corresponding vectors using the Amaxa Nucleofector Electroporation System. After one week of cell culture, cells were selected under 0.6 μg/ml Puromycin to enrich for electroporated cells and then sorted in single-cell wells using GFP as a marker, in order to generate a stable cell line.

### Hormone stimulation and SNAP labeling for SPT

16 hours before hormone stimulation (10nM R5020), the cell medium was changed to Phenol Red-free DMEM media supplemented with 10% charcoal-treated FBS Serum, 2mM L-glutamine, 1 mM sodium pyruvate, and 100 U/ml penicillin-streptomycin; from now on abbreviated as “charcoalized white DMEM”. We changed the medium as standard DMEM medium contains Phenol Red, which has interfering estrogenic activity with our experiments. The Janelia Fluor 549 dye coupled to the SNAP substrate was kindly provided by Luke Lavis (Janelia Farm, Ashburn, Virginia, USA). Cells were incubated with 10 nM for SPT in charcoalized white DMEM for 30 min at 37°C. Subsequently, the cells were washed three times with PBS, and then placed back in the incubator in charcoalized white DMEM for a one-hour washout at 37°C. This step was repeated two times. After the JF549 SNAP labeling, hormone stimulation was done using 10 nM R5020 (Promegestone) solubilized in Ethanol. The R5020 incubation time for SPT was 60 min.

### Western Blot

Cell lysates were resolved on SDS-polyacrylamide gels, and the proteins were transferred to Hybond-ECL nitrocellulose membranes (Amersham). Membranes were blocked with TBS-0.1% Tween20 (TBS-T) with 5% of skimmed milk, incubated for 1h at room temperature with a primary antibody (For antibody PR, sc-166169 from Santa Cruz Biotechnology, and antibody for tubulin, T9026 from Sigma, St. Louis, MO, USA, antibody for GFP, A-11122, Invitrogen), diluted in TBS-T with 2.5% skimmed milk. After three washes with TBS-T, membranes were incubated for 1h at room temperature with horseradish peroxidase-conjugated secondary antibodies (GE Healthcare, Chicago, IL, USA). Antibody binding was detected by chemiluminescence on an LAS-3000 image analyzer (Fuji PhotoFilm, Tokyo, Japan).

### Experimental setup for SPT

SPT was performed in a Nikon N-STORM 4.0 microscope system for localization-based super-resolution microscopy, equipped with a TIRF 100x, 1.49 NA objective (Nikon, CFI SR HP Apochromat TIRF 100XC Oil). The sample was illuminated by a continuous 561 nm laser line with a power of 30 mW before the objective in HILO configuration. The emission fluorescence was projected into an EM-CCD Andor Ixon Ultra Camera at a framerate of 15 ms. During imaging, the temperature was kept at 37°C by an incubation chamber.

### SPT data analysis

SPT trajectories were reconstructed as explained in our previous work (29). Briefly, single fluorophore localizations were detected using Trackmate (57) with sub-pixel localizations. The detected localizations were reconnected using the algorithm Linear Assignment Problem (LAP), and the minimum trajectory length for further analysis was 10 frames. To calculate the time-averaged MSD, we applied the following formula (29):

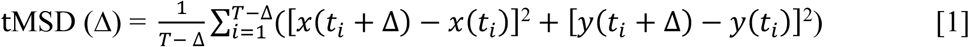

where *x* and y are the positions of the fluorophore, *t_i_* are the time steps, Δ is the time interval, and *T* the total duration of the trajectory. To extract the instantaneous diffusion coefficient (D_2-4_), we fit the 2^nd^ and 4^th^ point (Δ=2-4) of the MSD curve with the following formula:

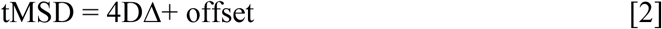

where D is the diffusion coefficient. For the D_2-4_ distributions, extracted D_2-4_ values below 0.0001 μm^2^/s were considered as fitting errors and discarded.

For the Spot-On Analysis, the MATLAB software version was downloaded from their dedicated website: https://github.com/elifesciences-publications/spot-on-matlab. The trajectories were fitted to a two-population kinetic model with the following parameters: dZ=0.7 μm, GapsAllowed=1, TimePoints=8, JumpsToConsider=4, ModelFit=2, NumberOfStates=2, D_Free_2State = [0.5;25]; D_Bound_2State = [0.0001;0.05].

### mRNA-sequencing samples, library preparation, and sequencing

RNA-sequencing experiments were performed as biological triplicate for the cells expressing WT-PR, βNTD and βDBD mutants before and after the 10nM R5020 treatment for 6 hours. RNA from WT-PR and βNTD and βDBD cells was extracted using the Quick-RNA kit (ZYMO Research; R1055) according to the manufactureŕs instructions. Purified RNA was analyzed on a Bioanalyzer using an RNA Pico assay chip. PolyA plus RNA mRNA libraries were prepared using TruSeq Stranded RNA Library Prep Kit (Illumina) and sequenced using Illumina HiSeq 2500.

### mRNA-sequencing data processing and analysis

Bulk mRNA-sequencing was performed on MCF7 cells expressing WT-PR, βNTD and βDBD mutants before and after R5020 treatment. Each condition was analyzed in triplicate, yielding a total of 18 FASTQ files. Quality control of raw sequencing reads was conducted using FastQC (version 0.72). Reads were aligned to the human reference genome (hg38) using HISAT2 (version 2.1.0) implemented on the Galaxy platform (version 7). Aligned reads were quantified using featureCounts (version 2.0.1), and differential gene expression analysis was performed with DESeq2 (version 2.11.40.6). Genes were considered significantly differentially expressed if they exhibited a Benjamini-Hochberg–adjusted p-value < 0.05 and an absolute log₂ fold change greater than 0.58. For data visualization and downstream analysis, the R packages tidyverse (version 2.0.0), scales (version 1.4.0), and ggplot2 (version 3.5.2) were used.

### Gene Ontology (GO) and Regulatory Gene Set Analysis

GO Annotation performed using the Gene Set Enrichment Analysis (GSEA, http://software.broadinstitute.org/gsea/index.jsp) database v5 (58, 59). The significant cut-off p-value and FDR q-value < 0.05. Plots were done with the use of Prism (GraphPad Prism 10.0.3 for MacOS) and R4.0.2.

### RNA extraction and RT-qPCR

Purified RNA from WT-PR, βNTD and βDBD expressing cells was used for reverse transcription of 1µg RNA, which was performed using an OneScript® RT Mix for qPCR with gDNAOut (OZYME; OZYA012-100) according to the manufactureŕs instructions. Complementary DNA was quantified by qPCR using ONEGreen FAST qPCR Premix (OZYME; OZYA008-1000) at the PCR analyzer Applied Biosystems (7500). For each gene product, relative RNA abundance was calculated using the standard curve method and expressed as relative RNA abundance after normalizing against the human GAPDH gene level as described (60, 61). All gene expression data generated by RT-qPCR represented the average and ± SEM of at least three biological replicates. Primers used for RT-qPCR are listed in **Table S1**.

### Cell Proliferation

Cell proliferation was performed using the BrdU (5’-bromo-2’-deoxyuridine) cell proliferation assay: WT-PR, βNTD and βDBD expressing cells (0.3 × 10^3^) were seeded in 96-well plates with complete DMEM media for 36 hours, incubated at 37 °C with 5% CO₂. Next, cells were washed with PBS and replaced with “charcoalized white DMEM” for 16 hours before R5020 induction for 24 hours. The cell proliferation ELISA BrdU (5’-bromo-2’-deoxyuridine) colorimetric assay (Roche,11647229001) was performed as per the manufacturer’s instructions. The experiments were performed on at least eight biological replicates.

### Cell Migration

Migration method was adopted from Justus et al. 2023 (62). WT-PR βNTD and βDBD expressing cells were seeded at a density of 2 × 10⁶ cells per P10 dish in complete DMEM medium and incubated at 37 °C with 5% CO₂. After 24 hours, the medium was removed, cells were washed with PBS, and the culture medium was replaced with “charcoalized white DMEM” before 10 nM R5020 induction for 24 hours. Cells were washed with pre-warmed PBS, detached, and washed again before being resuspended in migration buffer (DMEM supplemented with 10 mM HEPES and 0.1% BSA, pH 7.4). Cell viability was assessed, and the suspension was adjusted to a final concentration of 0.7 × 10⁶ viable cells/mL. Migration assays were conducted using 24-well Transwell inserts with 8 μm pores (Corning-Falcon, ref: 353097). A total of 600 μL of migration buffer containing 10% FBS as a chemoattractant was added to the lower chamber. Then, 100 μL of the prepared cell suspension (containing 1 × 10⁵ cells) was carefully added to the upper chamber. Plates were incubated overnight at 37 °C with 5% CO₂. The following day, non-migrated cells were gently removed from the upper surface of the membrane using a cotton swab. Migrated cells on the basal side were fixed in 4% paraformaldehyde and stained with crystal violet. Membranes were imaged under a light microscope at 10× and 20× magnifications. For quantitative analysis, 30% acetic acid was added to elute the dye, and the absorbance was measured at 565 nm using a CLARIOstar (BMG Labtech) plate reader.

### Machine Learning

The pipeline used to characterize the diffusive properties of the various constructs is mainly based on STEP (38). Given an input SPT trajectory, STEP uses state-of–the-art deep learning methods to predict the diffusive properties (diffusion coefficient and anomalous diffusion exponent) for every frame. This allows us to maximize the temporal resolution at which we analyze the trajectories, unveil changes of diffusion properties, and identify different diffusive states (63, 64). We post-process the frame-wise raw diffusive properties prediction with a kernel change detection algorithm to obtain segments of constant properties (65). This allows us to identify the diffusive changepoints as well as generally improve the analysis accuracy (38).

For this work, we have performed a series of adjustments over the original STEP architecture and have trained a new model using the tools provided by the authors (38). More precisely, we train a STEP model only with noiseless fractional Brownian motion trajectories, as opposed to using multiple anomalous diffusion methods with various noise levels. This ensures that the lowest diffusion coefficient predicted in the experiment matches the localization noise and removes all the prediction artifacts in trajectories with low diffusion coefficients. Additionally, we sample a random trajectory length between 20 and 200 for every batch that we use to take a random slice of all the trajectories therein. This makes the model robust to trajectory length variation while avoiding padding and keeping the training loop as efficient as possible. Finally, based on the findings of (63), we improve the original STEP by considering as input the motion displacements instead of using the raw coordinates. To do so, we compute the displacements for every dimension of an input trajectory and take the logarithm of their absolute value. Then, we compute the angle between consecutive displacements, setting the first one to zero by convention. We feed the STEP model this trajectory representation, which is rotationally invariant and preserves all the physically relevant information. To perform the analysis of the predictions, we first take all the trajectory segments for the three constructs, aggregate them in a dataset, and run a k-means clustering algorithm over it. We choose to do the clustering with all segments to avoid any potential bias some of the constructs may have, and improve the statistical relevance of the analysis with more data. We find that a separation into two clusters gives the best qualitative result.

To reveal changes in the different mutants’ distribution, we first constructed joint probability distributions of the diffusion coefficient 𝐷and anomalous exponent 𝛼for each of them by estimating normalized two-dimensional histograms in the (𝐷, 𝛼) plane. These histograms were converted into discrete probability masses by accounting for the bin areas. Pairwise differences between conditions were then quantified using a symmetric, normalized contrast,

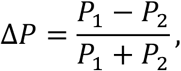

computed pointwise over the (𝐷, 𝛼)space. This definition yields a bounded measure [−1,1] that highlights relative enrichments or depletions between conditions while suppressing regions of low overall probability. The resulting difference maps provide a direct visualization of how the distributions shift across dynamical regimes.

### Quantification and statistical analysis

For RT-qPCR experiments, a Two-tailed unpaired Student’s t-test was used to determine statistical significance between the groups. For cell proliferation and migration experiments, the significance between groups was calculated by the Wilcoxon-Mann-Whitney test. Plots and indicated statistical analysis were done with the use of Prism (GraphPad Prism 10.0.3 for MacOS) unless otherwise stated. If exact p-values are not shown or indicated in the legend, then p-values are represented in all figures as follows: *, p-value ≤ 0.05; **, p-value ≤ 0.01; ***, p-value ≤ 0.001; °, p-value > 0.05.

## Acknowledgments

We thank Gordon Hager (NIH) for providing the pGFP-PRB plasmid and Luke Lavis (Janelia Research Campus) for kindly providing the SNAP-Janelia Fluor 549 dye. We would like to thank the Advanced Light Microscopy Unit, the Genomics facility, and the Flow Cytometry Unit of the Center for Genomic Regulation (CRG, Barcelona) for their support. The research leading to these results has received the following funding: J.A.T.-P. and C.R. jointly from BIST-Ignite funding (PHASE-CHROM). J.A.T.-P. from the Government of Spain JdC-IJCI-2017-33160 and RYC2023-044870-I. M.G.-P. from the European Commission ERC Adv788546 (NANO-MEMEC), the Government of Spain (Severo Ochoa CEX2019-000910-S and PID2023-147711NB-I00), Fundació CELLEX (Barcelona), Fundació Mir-Puig, and the Generalitat de Catalunya through the CERCA program and AGAUR (Grant No. 2021 SGR 01450).

B.R. and M.L. acknowledge support from MCIN/AEI (PGC2018-0910.13039/501100011033, CEX2019-000910-S/10.13039/501100011033, Plan Nacional STAMEENA PID2022-139099NB, project funded MCIN and by the “European Union NextGenerationEU/PRTR" (PRTR-C17.I1), FPI); Ministry for Digital Transformation and of Civil Service of the Spanish Government through the QUANTUM ENIA project call - Quantum Spain project, and by the European Union through the Recovery, Transformation and Resilience Plan - NextGenerationEU within the framework of the Digital Spain 2026 Agenda; CEX2024-001490-S [MICIU/AEI/10.13039/501100011033]; Fundació Cellex; Fundació Mir-Puig; Generalitat de Catalunya (European Social Fund FEDER and CERCA program); Barcelona Supercomputing Center MareNostrum (FI-2023-3-0024); Funded by the European Union (HORIZON-CL4-2022-QUANTUM-02-SGA, PASQuanS2.1, 101113690, EU Horizon 2020 FET-OPEN OPTOlogic, Grant No 899794, QU-ATTO, 101168628), EU Horizon Europe Program (No 101080086 NeQSTGrant Agreement 101080086 — NeQST).). C.G-C. and X.S. were supported by AGAUR (2021 SGR 476), AEI (PID2022-141816OB-I00), and the ERC (CONCERT, contract number 648201). G.M.-G. acknowledges support from the European Union (ERC Advanced Grant, QuantAI, No. 101055129). P.S. received funding from the French National Research Agency (ANR) Young Investigator grant (ANR-21-CE12-0010), Cancer Research Foundation (ARCPJA22021050003683), La Ligue Foundation for Cancer (Haute-Garonne, 285458), Centre national de la recherche scientifique (CNRS), Université de Toulouse, Toulouse, France. The views and opinions expressed in this article are, however, those of the author(s) only and do not necessarily reflect those of the European Union or the European Research Council - neither the European Union nor the granting authority can be held responsible for them.

## Author Contributions

G.M.-G., M.L., M.F.G.-P., J.A.T.-P., P.S. supervised research; C.R.-A., J.A.T.-P., M.F.G.-P., P.S. designed research; L.R., D.S., C.R.-A., J.A.T.-P., L.E., L.I.L.C., S.N., C.G.-A., E.S., N.Y., P.S. performed research; L.R., B.R., G.M.-G., C.G.-A., E.S., J.A.T. N.Y., D.S., R.L.K., P.S. analyzed data; L.R., X.S., C.G.-C., M.L., G.M.-G., M.F.G.-P., P.S. and J.A.T.-P. wrote the paper.

## Declaration of Interests

X.S. is founder and scientific advisor of Nuage Therapeutics.

## Resource Availability

RNA sequencing data performed and used in this study have been deposited at GEO under the accession code GSE308408 (secure token to access is ojilyckqtrujjuz) and will be publicly available on acceptance for publication. Further information and requests for resources and results should be directed to the corresponding authors, Maria Garcia-Parajo, Juan A. Torreno-Pina, and Priyanka Sharma.

---

**Supplementary Figure 1.**
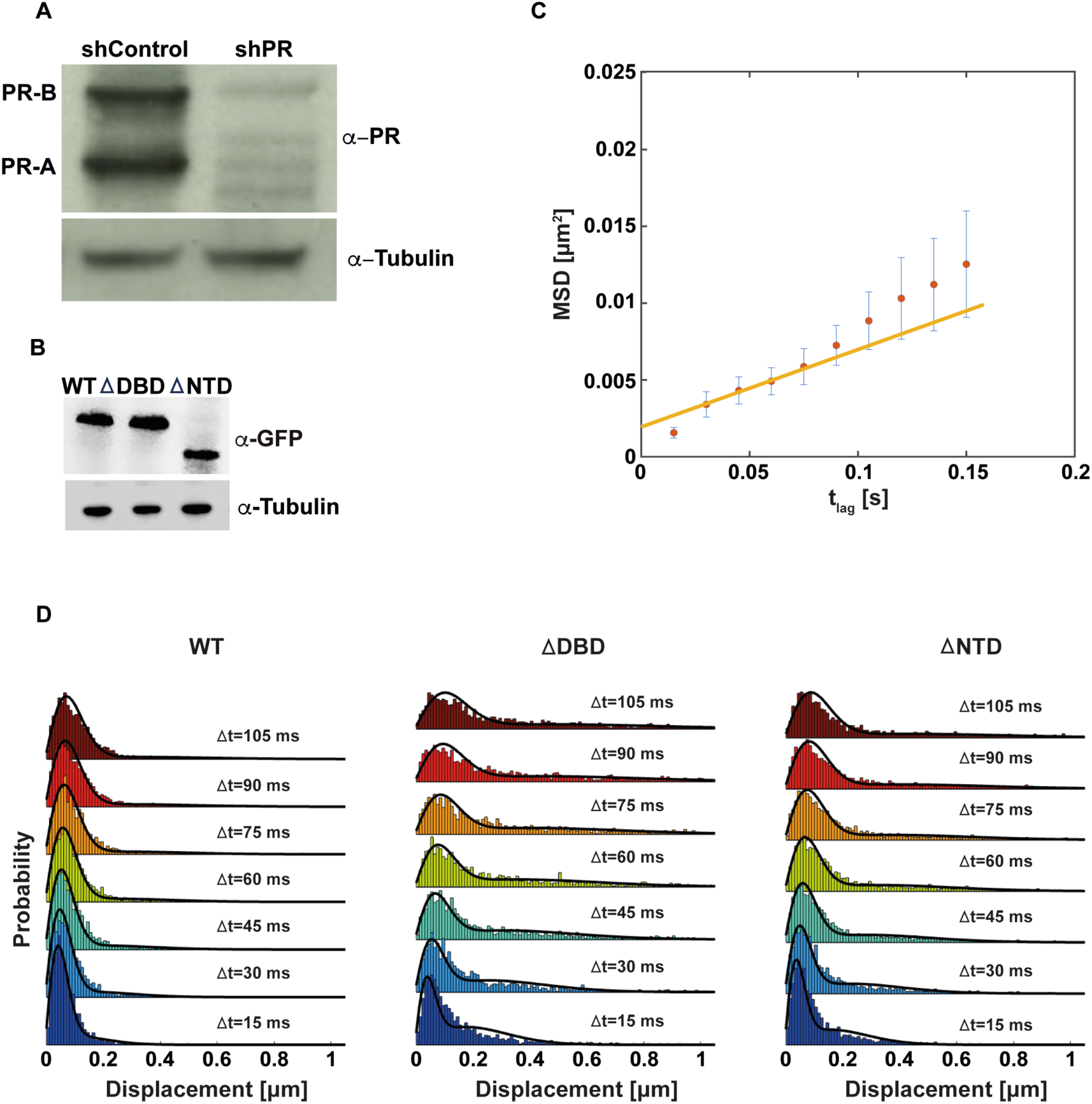
: **SPT analysis.** (**A**) Extract from shControl and shPR showing the effective knockdown of endogenous PR as indicated by western blot for the indicated antibodies. (**B**) Extracts from WT, βDBD, or βNTD cells followed by the western blot of the indicated antibodies showing the knock-in expression of PR. (**C**) Representative MSD plot of an individual SPT trajectory. The MSD and the errors are represented as red dots and blue error lines, respectively. The fitting through the 2^nd^-4^th^ points of the MSD plot is represented as a yellow line (see Methods). (**D**) Representative histograms of the displacements in dependence on time lags (insets) as extracted from Spot-On analysis (see Methods) of WT, βDBD, or βNTD. In the case of βDBD and βNTD, the distribution of the displacements is shifted towards longer displacements.

**Supplementary Figure 2.**
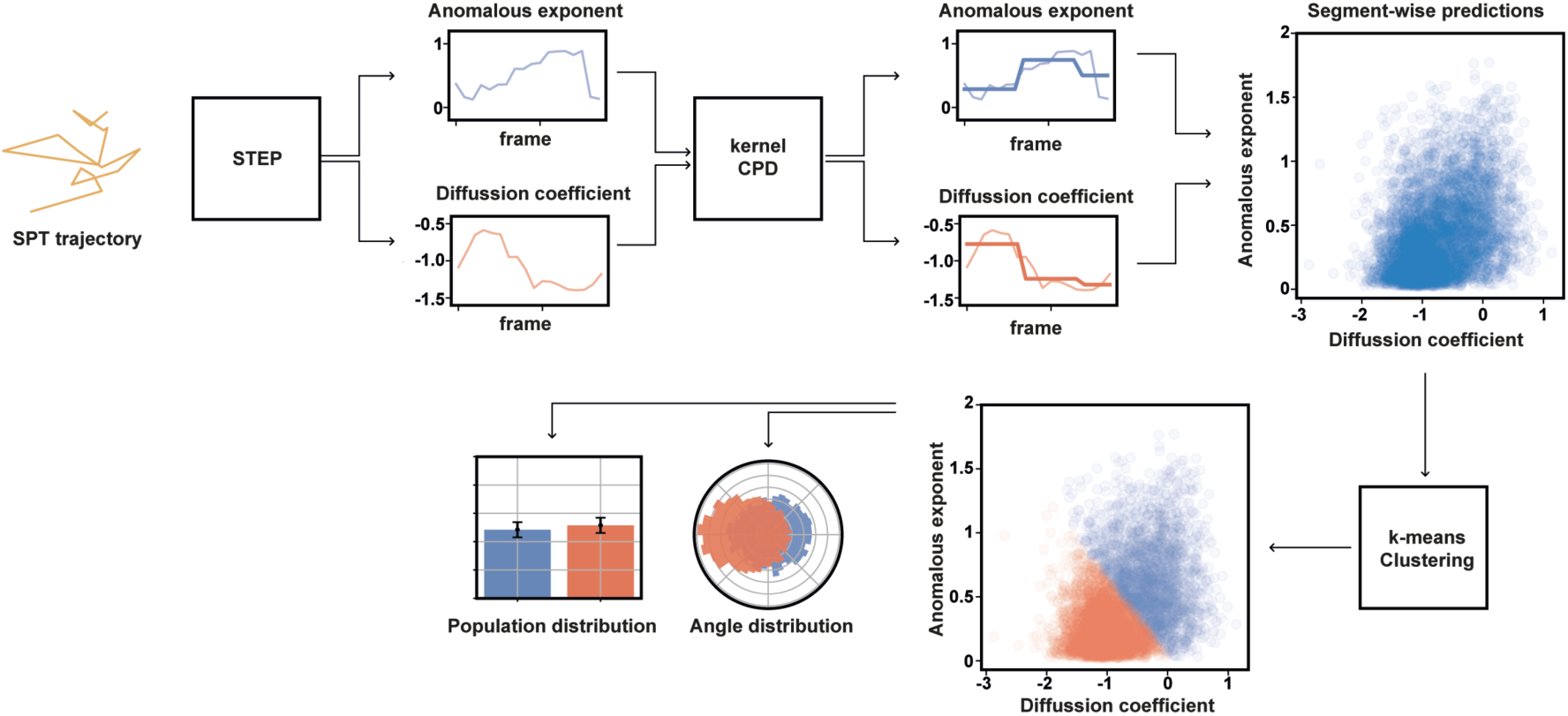
: Deep learning pipeline. Given a trajectory, STEP predicts diffusive properties for each frame. We then use a kernel changepoint detection algorithm to segment these predictions. From the obtained segment, we perform clustering via k-means. The resulting clusters are then used to compute population and angle distribution as well as the confinement radius of the bound state.

**Supplementary Figure 3.**
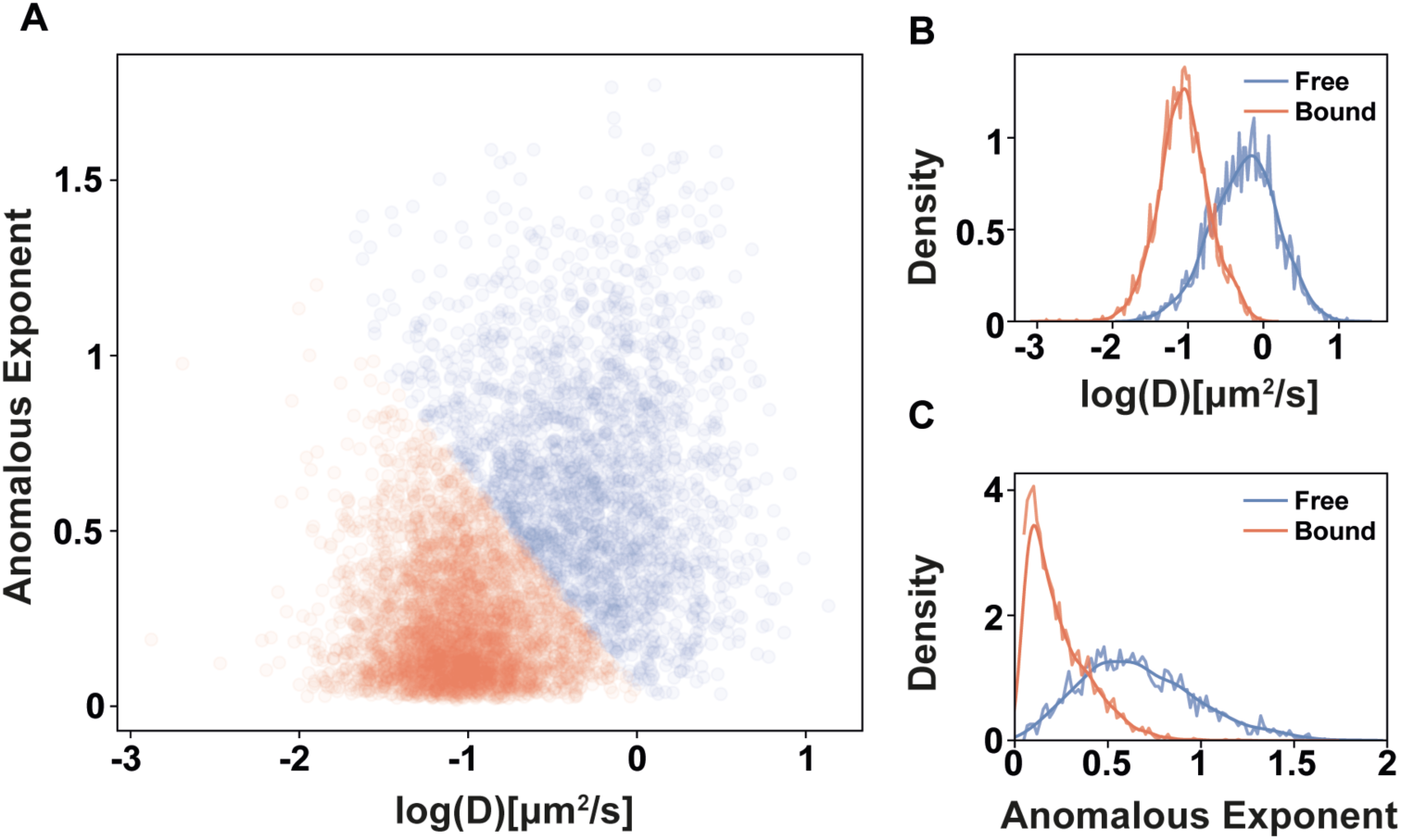
: K-means clustering of trajectory segments. (**A**) Clustering of the segments as predicted by k-means with two target clusters. (**B**, **C**) Diffusion coefficient and anomalous exponent distribution of the two found clusters. The actual distribution and a kernel density estimation are represented.

**Supplementary Figure 4.**
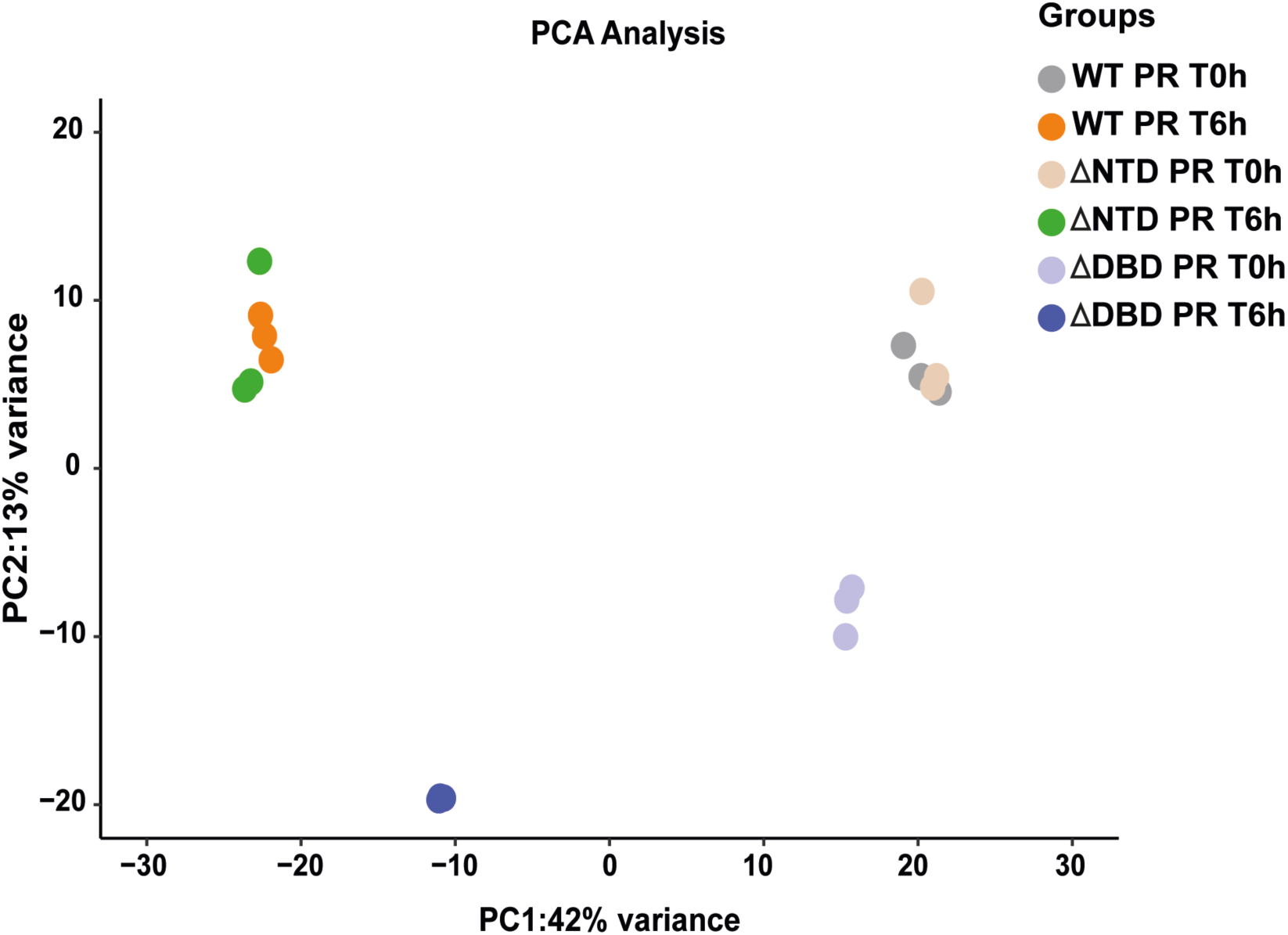
: Principal Component Analysis (PCA) of transcriptional profiles across experimental groups. PCA plot illustrates the variance in transcriptional profiles among six experimental groups of PR expressing cells with R5020 induction: WT (T0), WT (T6h), βNTD (T0), βNTD (T6h), ΔDBD (T0), and βDBD (T6h). PC1 accounts for 42% of the variance, and PC2 accounts for 13%. Each dot represents a biological triplicate, color-coded based on the group as indicated. The separation of clusters emphasizes the transcriptional impact of the WT-PR and PR mutants (βNTD or βDBD) in response to R5020 induction.

**Supplementary Figure 5.**
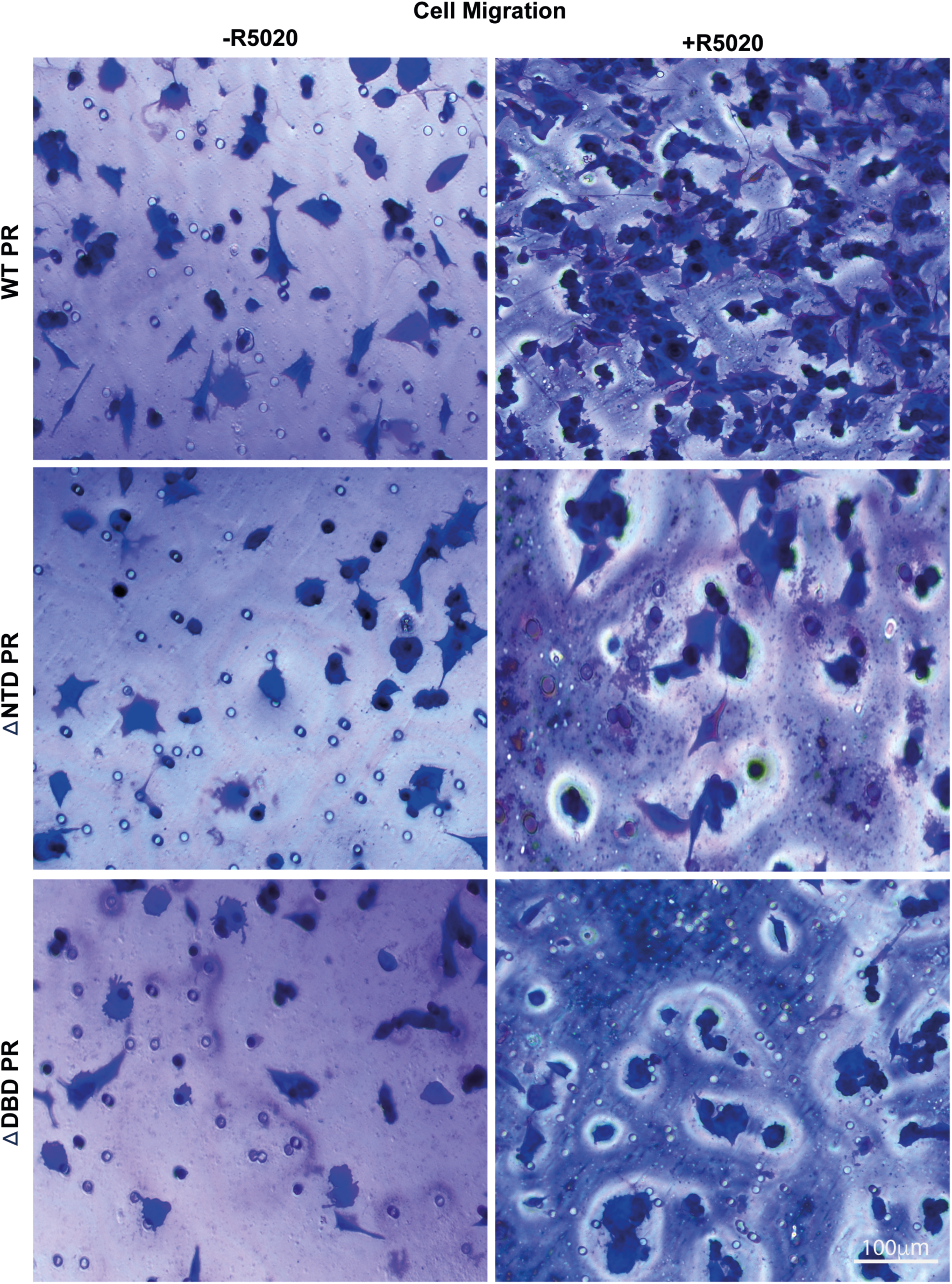
: Cell Migration. Representative Crystal Violet staining microscopy images of MCF7 cells expressing WT-PR and PR mutant (βNTD or βDBD) before (-R5020) and after (+ R5020) induction treatment during the transwell cell migration assay. Scale bar 100μm.

**Supplementary Table S1.**
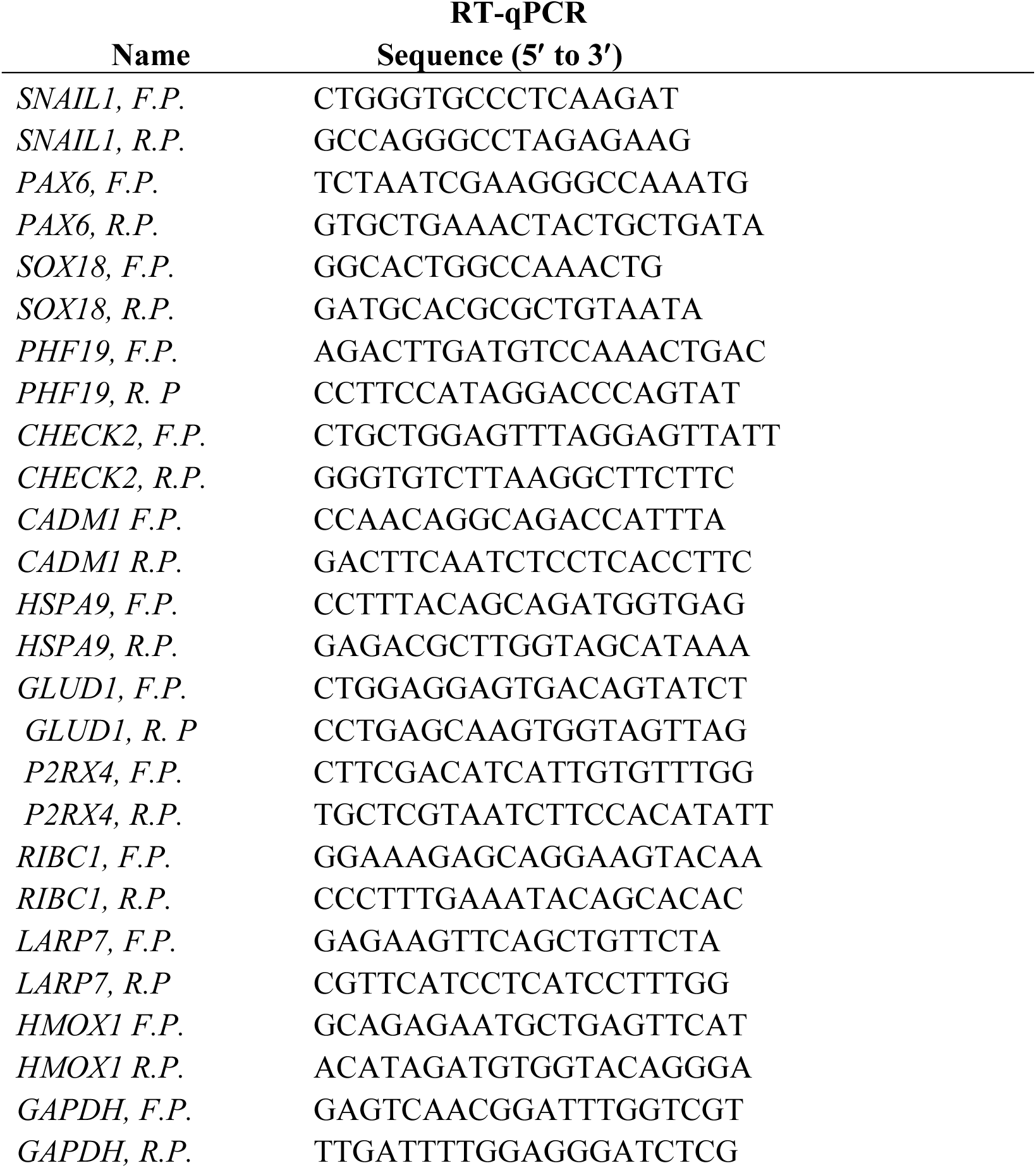
- Primers used in this study.

## Notes

### Competing Interest Statement

Xavier Salvatella is founder and scientific advisor of Nuage Therapeutics.

